# Anabolic sensitivity in healthy, lean, older men is associated with higher expression of amino acid sensors and mTORC1 activators

**DOI:** 10.1101/2024.06.15.598989

**Authors:** Oscar Horwath, Marcus Moberg, Nathan Hodson, Sebastian Edman, Mats Johansson, Eva Andersson, Gerrit van Hall, Olav Rooyackers, Andrew Philp, William Apró

**Affiliations:** Department of Physiology, Nutrition and Biomechanics, the Swedish School of Sport and Health Sciences, Stockholm, Sweden; Department of Physiology and Pharmacology, Karolinska Institute, Stockholm, Sweden; Department of Women’s and Children’s Health, Karolinska Institute, Stockholm, Sweden; Department of Exercise Sciences, Faculty of Kinesiology and Physical Education, University of Toronto, Toronto, Ontario, Canada; Department of Sport and Exercise Sciences, Institute of Sport, Manchester Metropolitan University, Manchester, UK; Division of Clinical Chemistry, Department of Laboratory Medicine, Karolinska Institute, Stockholm, Sweden; Department of Molecular Medicine and Surgery, Karolinska Institute, Stockholm, Sweden; Department of Biomedical Sciences, Faculty of Health and Medical Sciences, University of Copenhagen, Copenhagen, Denmark; Clinical Metabolomics Core Facility, Department of Clinical Biochemistry, Rigshospitalet, University of Copenhagen, Copenhagen, Denmark; Department of Clinical Science, Intervention and Technology, Karolinska Institute, Stockholm, Sweden; Centre for Healthy Ageing, Centenary Institute, Sydney, NSW, Australia; School of Sport, Exercise and Rehabilitation Sciences, University of Technology Sydney, NSW, Australia

**Keywords:** sarcopenia, protein synthesis, cell signaling, resistance exercise, amino acid sensing

## Abstract

**Background:** Sarcopenia is thought to be underlined by age-associated anabolic resistance and dysregulation of intracellular signalling pathways. However, it is unclear whether these phenomena are driven by ageing *per se* or other confounding factors.

**Methods:** Lean and healthy young (n=10, 22 ± 3 yrs, BMI; 23.4 ± 0.8 kg/m^2^) and old men (n=10, 70 ± 3 yrs, BMI; 22.7 ± 1.3 kg/m^2^) performed unilateral resistance exercise followed by intake of essential amino acids (EAA). Muscle biopsies were collected from the rested and the exercised leg before, immediately after, as well as 60 and 180 minutes after EAA intake. Muscle samples were analyzed for amino acid concentrations, muscle protein synthesis (MPS) and associated anabolic signaling.

**Results:** Following exercise, peak plasma levels of EAA and leucine were similar between groups, but the area under the curve was ∼11% and ∼28% lower in Young (p<0.01). Absolute levels of muscle EAA and leucine peaked 60 min after exercise, with ∼15 and ∼21 % higher concentrations in the exercising leg (p<0.01) but with no difference between groups. MPS increased in both the resting (∼ 0.035%·h^-1^ to 0.056%·h^-1^, p<0.05) and exercising leg (∼ 0.035%·h^-1^ to 0.083%·h^-1^, p<0.05) with no difference between groups. Phosphorylation of S6K1^Thr389^ increased to a similar extent in the exercising leg in both groups but was 2.8-fold higher in the resting leg of Old at the 60 min timepoint (p<0.001). Phosphorylation of 4E-BP1^Ser65^ increased following EAA intake and exercise, but differences between legs were statistically different only at 180 min (p<0.001). However, phosphorylation of this site was on average 78% greater across all timepoints in Old (p<0.01). Phosphorylation of eEF2^Thr56^ was reduced (∼ 66 and 39%) in the exercising leg at both timepoints after EAA intake and exercise, with no group differences (p<0.05). However, phosphorylation at this site was reduced by ∼ 27% also in the resting leg at 60 min, an effect that was only seen in Old (p<0.01). Total levels of Rheb (∼ 45%), LAT1 (∼ 31%) and Rag B (∼ 31%) were higher in Old (p<0.001).

**Conclusion:** Lean and healthy old men do not manifest AR as evidenced by potent increases in MPS and mTORC1 signalling following EAA intake and exercise. Maintained anabolic sensitivity with age appears to be a function of a compensatory increase in basal levels of proteins involved in anabolic signalling. Therefore, our results suggest that age *per se* does not appear to cause AR in human skeletal muscle.

## Introduction

Age-associated muscle loss, i.e. sarcopenia, is a debilitating condition that leads to several negative health outcomes and in some cases, premature mortality. [1, 2]. The current prevailing view is that sarcopenia is driven primarily by anabolic resistance (AR) [3, 4]. As a phenomenon, AR is defined as a reduced ability of aged muscle to increase muscle protein synthesis (MPS) in response to anabolic stimuli such as essential amino acids and/or resistance exercise.

At the molecular level, MPS is largely regulated by the mechanistic target of rapamycin (mTORC1) pathway [5, 6] with mTORC1 inhibition shown to block both EAA and resistance exercise-induced increases in MPS [7, 8]. *In-vitro* studies on transformed cell lines have identified the lysosome as a key intracellular compartment where mTORC1 is activated by amino acids [9, 10].

During amino acid sufficiency, mTORC1 is translocated by Rag GTPases, from the cytoplasm to the lysosome, where it becomes fully activated by Rheb (GTP-bound ras-homolog enriched in brain) [9, 10]. The importance of the Rags in this process is illustrated by the complete inactivation of mTORC1 by genetic knock-down of the Rag proteins [9]. As such, dysregulation of mTORC1 complex assembly may be, at least in part, an underlying cause of AR. In human skeletal muscle, this notion is supported by blunted mTORC1 activity in older adults, despite an ample supply of amino acids [11, 12].

While AR is an established physiological phenomenon, it is important to note that not all studies support the existence of AR in aging populations [13]. These contradictory findings suggest that confounding factors other than age *per se* may be involved in the development of AR. These include increased adiposity and inactivity, both of which have been shown to blunt MPS in response to protein feeding [14–17]. Furthermore, studies examining the upstream regulation of the mTORC1 pathway in human skeletal muscle are scarce and with conflicting results [18–20]. Importantly, no previous studies have examined the regulation of MPS in healthy, lean, physically active older men or effectors such as Rheb and Rags in aged human skeletal muscle.

Thus, to gain further insights into the underlying mechanisms of AR, and the role of age *per se*, the primary aim of this study was to investigate the anabolic response in lean, healthy, and physically active young and old men. Participants performed unilateral resistance exercise followed by consumption of essential amino acids, and muscle samples were analyzed for MPS and mTORC1 signaling. Muscle samples were also analyzed for lysosomal mTORC1 interaction as well as Rheb and Rag protein content. It was hypothesized that ageing would lead to AR as evident by blunted MPS and mTORC1 signaling following EAA ingestion and resistance exercise.

## Methods

### Ethical approval

Participant recruitment and study experiments were conducted at the University of Birmingham, Birmingham UK (Young) and the Swedish School of Sport and Health Sciences in Stockholm, Sweden (Old). The study was approved by ethical review boards at both research locations (West Midlands, Black Country Research Ethics Committee, #17/WM/0068 and the Swedish Ethical Review Authority, #2017/2107-31/2) and performed in accordance with the Declaration of Helsinki. All participants provided written informed consent prior to participating in the study.

### Participants

Ten young (Young; 18-35 yrs) and ten old (Old; 65-75 yrs) men were recruited for this study. All participants were required to be non-smoking, lean (body mass index; BMI < 25) and in general good health without any cardiovascular or metabolic diseases. They were not allowed to consume any medications with known side effects related to skeletal muscle metabolism. None of the participants in the Young group consumed any medications. In the Old group, three participants consumed medication against acid reflux (n=1, Esomeprazole), hypertension (n=1, Amlodipine and Losartan) and against benign prostate hyperplasia (n=1, Finasteride). Participants were required to be recreationally active 2-3 times per week but were not allowed to be involved in regular structured resistance exercise. Participant characteristics were as follows (mean ± SD) for Young vs Old: age; 22 ± 3 vs 70 ± 3 yrs, weight; 74 ± 5 vs 73 ± 6 kgs, height; 177 ± 5 vs 179 ± 5 cm, BMI; 23.4 ± 0.8 vs 22.7 ± 1.3 kg/m2, leg strength; 58 ± 6 vs 30 ± 7 kg.

### Study design

Participants visited the laboratory in the morning on four different occasions, each time in a fasted state, having refrained from vigorous physical activity for the last 48 hours. Each visit was separated by 5-7 days. On the first visit, leg strength was determined in a leg-extension machine as the 10-repetition maximum (10RM) for the leg previously randomized to perform the resistance exercise. During the second and third visits, participants performed two familiarization sessions during which they performed the exercise protocol based on the previously determined 10RM. The purpose of the familiarization sessions was to ensure proper technique, adjust the load if needed, and to minimize the risk of a general stress response to the exercise during the experimental trial.

On the day of the experimental trial (Visit 4), participants reported to the laboratory at ∼ 7.00 AM (Figure 1). Upon arrival, they were placed in a supine position and peripheral venous catheters were inserted in the antecubital vein of each arm for repeated blood sampling and tracer infusion. Following a 30 min resting period, a baseline blood sample was drawn after which a primed constant infusion of L-[*ring*-^13^C_6_]-phenylalanine (0.05 µmol · kg ^-1^ · min ^-1^, prime 2 mol ·kg ^-1^; Cambridge Isotope Laboratories, Danvers, MA) was initiated and maintained for the duration of the entire experiment (∼ 7 h). Participants remained resting in the supine position for 150 min after which local anesthesia was administered and a baseline biopsy collected from the *m. vastus lateralis* of the resting leg using a Bergström needle with manually applied section. Next, the unilateral resistance exercise protocol was initiated with three warm-up sets with 10 repetitions at approximately 30, 50 and 70% of 10 RM, with two minutes of rest between each set. Following warm-up, participants performed 10 sets of 10 repetitions with three-minute rest-intervals between each set. Each participant started at their individual 10 RM, and the load was gradually lowered when proper form could no longer be maintained for 10 repetitions. Immediately after completion of the exercise protocol, participants were instructed to consume an essential amino acid drink (240 mg EAA/kg bw; Ajinomoto, Kanagawa, Japan) as fast as possible (< 1 min). The EAA composition of the drink was as follows: 17% leucine, 9% isoleucine, 11% valine, 14% histidine, 18% lysine, 3% methionine, 14% phenylalanine and 14% threonine. To maintain steady state plasma enrichment, the EAA drink also included 4% L-[ring-^13^C_6_]-phenylalanine. During recovery, additional muscle biopsies were collected in both the exercising and resting leg at 60 and 180 min after EAA intake. For each sampling, a new incision was made, and the first biopsy was taken approximately 8–9 cm above the mid patella and the following biopsies approximately 3–4 cm proximal to the previous one. Biopsy samples were immediately blotted free of blood and frozen in liquid nitrogen and stored at -80°C for later analysis.

**Figure 1.**
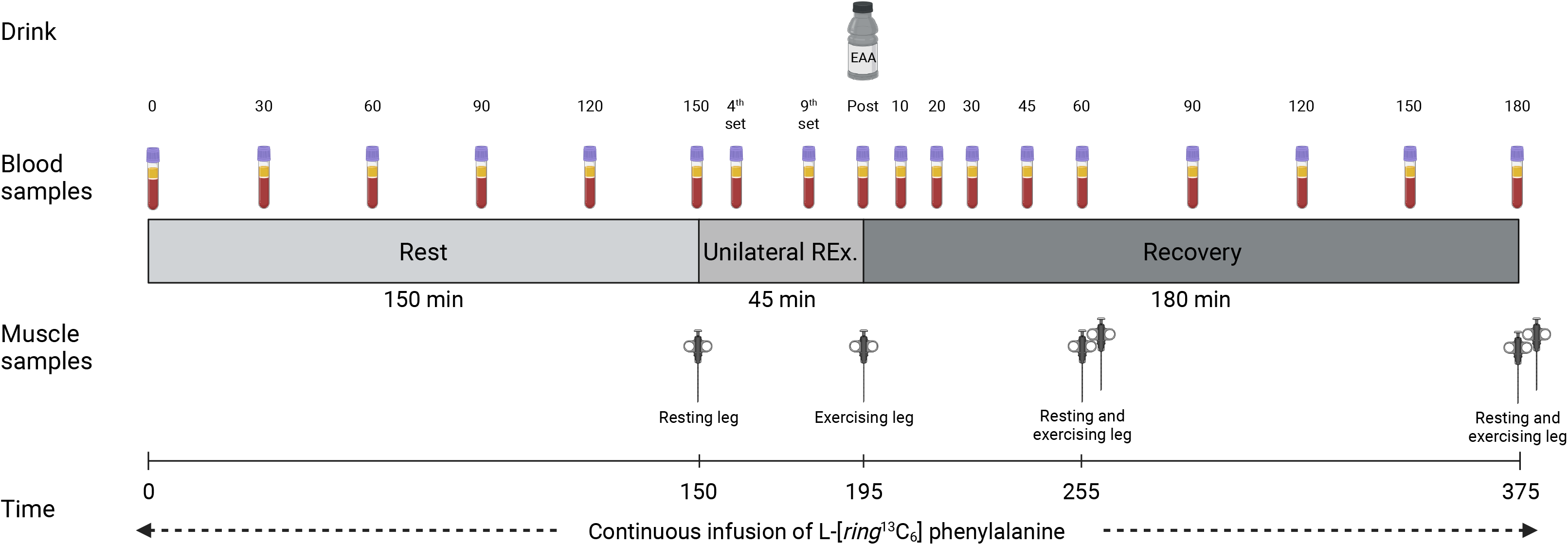
Schematic overview of the experimental trial. This figure was created with BioRender.com.

Blood samples for analyses of insulin, lactate and amino acid concentrations, as well as plasma isotope enrichment were collected in EDTA-tubes throughout the entire experimental trial. All blood samples were kept on ice until the end of the experimental trial at which they were centrifuged for 10 minutes at 3000 g. The plasma obtained was stored at -80°C for future analyses.

### Sample processing and analyses

Muscle samples were lyophilized, thoroughly cleaned, homogenized, and subsequently separated into three fractions (myofibrillar, lysosome enriched and cytosolic) as described in detail previously [21]. The myofibrillar fraction was processed and analyzed for isotopic enrichment of L-[ring-^13^C_6_]-phenylalanine for the calculation of myofibrillar protein synthesis rates (FSR) [22]. The lysosomal and cytosolic fractions were combined with Laemmli sample buffer (Bio-Rad Laboratories), and immunoblotting was performed as described previously [23]. Representative bands for each of the proteins are presented in Supplemental figure 3. Antibody information is presented in Table 1.

**Table 1.**
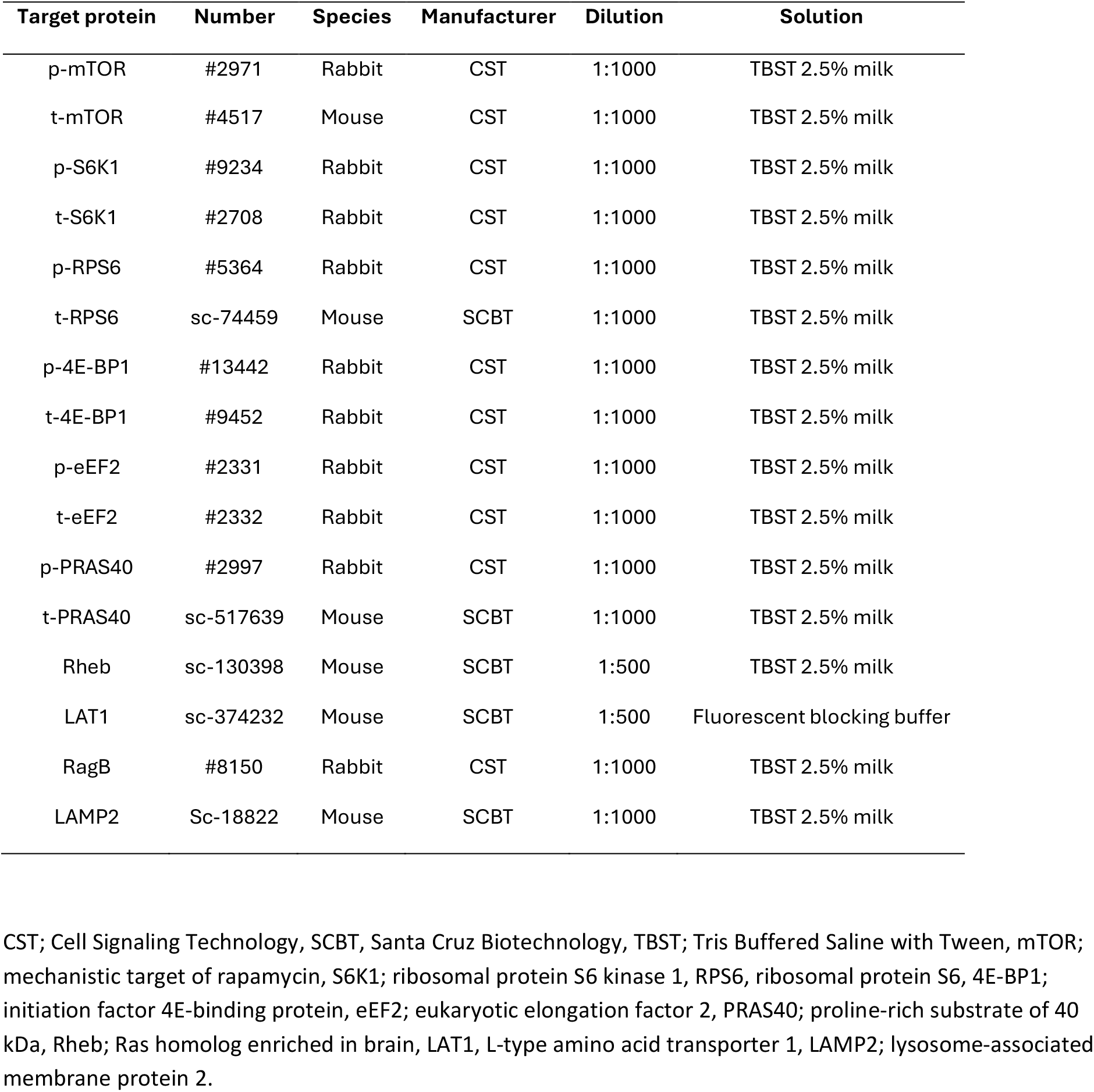
Antibody information.

For analysis of muscle amino acid concentrations, an aliquot of the cytosolic fraction was combined with equal volumes of ice cold trichloroacetic acid (5% v/v) and left on ice for 30 min, followed by centrifugation at 16000 g for 10 minutes at 4°C. The deproteinized supernatant was derivatized using the AccQ-Tag Ultra Derivatization Kit (Waters Sverige AB, Solna, Sweden) and subsequently analyzed using liquid chromatography-tandem mass-spectrometry (Xevo™ TQ MS, Waters). Free intracellular enrichment was analyzed on a separate aliquot of muscle (2-3 mg) as described previously [24]. Plasma lactate and insulin were analyzed spectrophotometrically as described previously [23].

### Statistical analysis

Data are presented as means ± standard deviation (SD) unless stated otherwise. Statistica 14 (TIBCO Software Inc, CA, USA) and GraphPad Prism version 10 (GraphPad Software Inc, USA) were used for analysis and data presentation. Data were initially checked for normality and skewed variables were log-transformed to be considered acceptable for parametric statistical testing. Plasma variables, i.e., lactate, insulin, and amino acid concentrations, were analyzed using a two-way (age x time) repeated-measures analysis of variance (ANOVA). The area under the curve (AUC) for the two groups was compared using two-tailed unpaired t-tests. Muscle amino acid concentrations, intracellular signaling, and FSR were analyzed using a three-way (age x time x leg) repeated-measures ANOVA. In this analysis, due to the unilateral study design, the sample obtained from the resting leg served as the basal value also for the exercising leg. For the same reason (unequal timepoints), the immediate exercise effect (pre-to-post) on intracellular signaling was analyzed separately using a two-way (age x time) ANOVA. Significant main or interaction effects were further analyzed with Bonferroni *post hoc* tests to locate differences. Statistical significance was set to p<0.05.

## Results

### Plasma lactate and insulin concentrations

Plasma lactate increased during exercise, peaked immediately after, and then returned to baseline levels during recovery (p<0.001 for time, Figure 2A). Peak lactate levels (8.2 and 3.5 mmol·L^-1^) and the AUC were ∼ 2.4-fold and 1.7-fold higher in Young than Old, respectively (p<0.001 for both). Plasma insulin increased shortly after EAA intake, peaked at ∼ 18 mU·L^-1^, but returned to baseline levels 90 min later, with no group differences (p<0.001 for time, Figure 2B).

**Figure 2.**
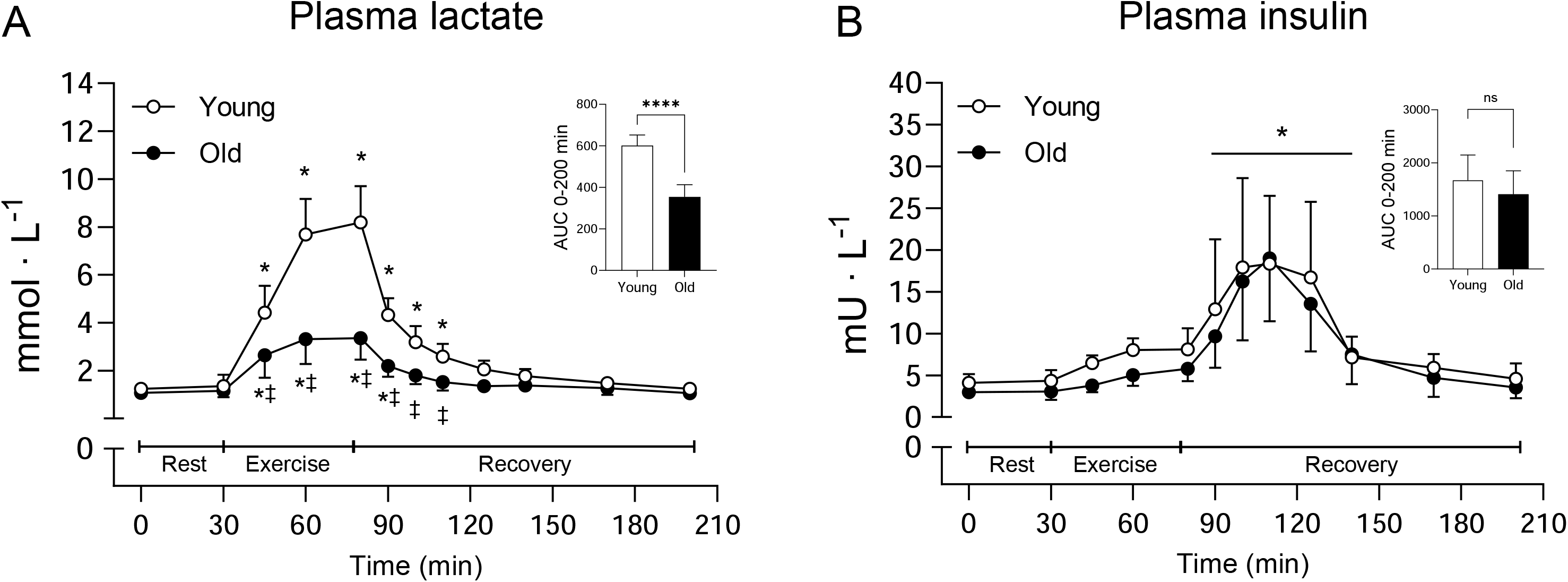
Plasma lactate (A) and insulin concentrations (B) before, during, and after resistance exercise and intake of EAA in Young (white circles) and Old (black circles). A two-way (age x time) repeated-measures ANOVA was used to analyze plasma concentrations. The ANOVA revealed a significant interaction (age x time) for (A) and a main effect of time for (B). The symbols marked without lines represents a three-way interaction, and symbols marked with a long line represents a main effect; * p<0.05 different from baseline, ‡ p<0.05 different from Young. The AUC was calculated for the entire time period (0-200 min). A two-tailed unpaired t-test was used to analyze AUC and symbols indicate; **** p<0.0001 different from Young. Values are presented as means ± SD for 20 participants. AUC; area under the curve, ns; not significant.

### Plasma and muscle amino acid concentrations

Plasma EAA increased following intake and remained elevated during the entire recovery period. Values were however numerically higher in Old compared to Young, resulting in a significantly larger AUC (p=0.007, Figure 3A). This was highly similar to the pattern observed for plasma BCAA and leucine alone where Old had ∼ 22% and 28% larger AUC compared to Young (p<0.001 for both, Figure 3C and 4A). The remaining EAAs (EAA-BCAA) and phenylalanine were also elevated during recovery, but no differences were observed between groups (p<0.001 for time, Figure 3E and 4C). Plasma tryptophan was ∼ 16% lower at rest in Old compared to Young but levels were reduced below baseline in a similar fashion for both groups in response to EAA intake and exercise (p<0.05 for both initial timepoints, Figure 4E).

**Figure 3.**
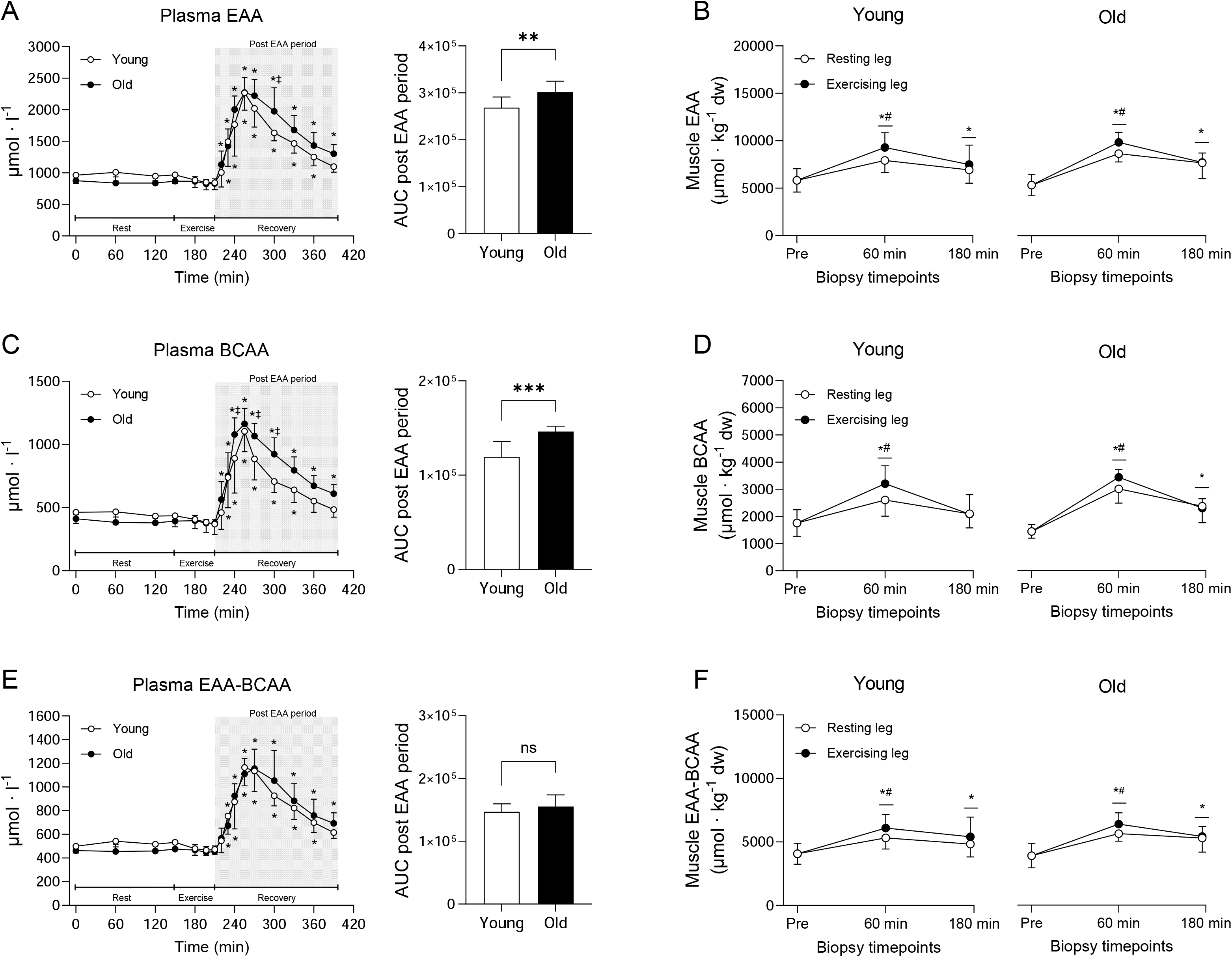
Plasma and muscle concentrations of EAA (A, B), BCAA (C, D) and the remaining essential amino acids (EAA-BCAA) (E, F) before, during, and after resistance exercise and intake of EAA in Young and Old. In plasma, Young are represented by white circles and Old are represented by black circles, whereas in muscle, white and black circles indicate the resting and the exercising leg, respectively. A two-way (age x time) repeated-measures ANOVA was used to analyze plasma concentrations. The ANOVA revealed a significant interaction (age x time) for all variables. The symbols marked without lines represent a three-way interaction and symbols marked with short lines represent a two-way interaction; * p<0.05 different from baseline, ‡ p<0.05 different from Young. The AUC was calculated for the entire post EAA period (180 min). A two-tailed unpaired t-test was used to analyze AUC and symbols indicate; ** p<0.01 and *** p<0.001 different from Young. A three-way (age x time x leg) repeated-measures ANOVA was used to analyze muscle concentrations. The ANOVA revealed a significant interaction (time x leg) for all variables. The symbols indicate; * p<0.05 different from baseline, # p<0.05 different from the resting leg. Values are presented as means ± SD for 20 participants. AUC; area under the curve, ns; not significant.

**Figure 4.**
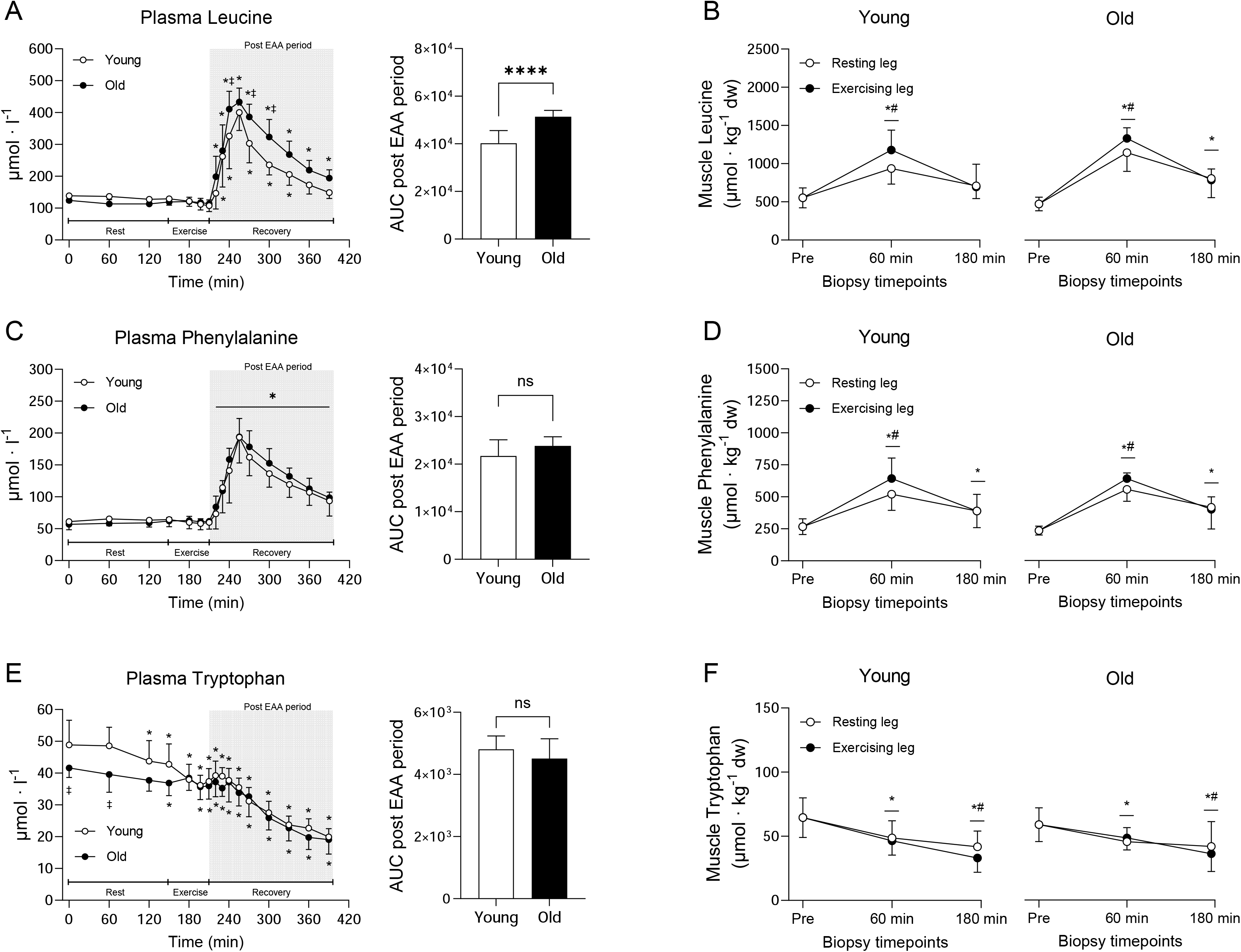
Plasma and muscle concentrations of leucine (A, B), phenylalanine (C, D) and tryptophan (E, F) before, during, and after resistance exercise and intake of EAA in Young and Old. In plasma, Young are represented by white circles and Old are represented by black circles, whereas in muscle, white and black circles indicate the resting and the exercising leg, respectively. A two-way (age x time) repeated-measures ANOVA was used to analyze plasma concentrations. The ANOVA revealed a significant interaction (age x time) for (A and E) and a main effect of time for (C). The symbols marked without lines represent a three-way interaction, symbols marked with short lines represent a two-way interaction and symbols marked with a long line represents a main effect; * p<0.05 different from baseline, ‡ p<0.05 different from Young. The AUC was calculated for the entire post EAA period (180 min). A two-tailed unpaired t-test was used to analyze AUC and symbols indicate; **** p<0.0001 different from Young. A three-way (age x time x leg) repeated-measures ANOVA was used to analyze muscle concentrations. The ANOVA revealed a significant interaction (time x leg) for all variables. The symbols indicate; * p<0.05 different from baseline, # p<0.05 different from the resting leg. Values are presented as means ± SD for 20 participants. AUC; area under the curve, ns; not significant.

Muscle EAA increased after intake and remained elevated throughout recovery, with ∼ 15% higher levels in the exercising leg compared to the resting leg at 60 min (p<0.01, Figure 3B). While there were no significant differences for absolute values, the fold change was significantly larger in Old compared to Young (p=0.007 for age, supplemental Figure 1E). This pattern was similar for muscle BCAA (Figure 3D and supplemental Figure 1D) and leucine alone (Figure 4B and supplemental Figure 1A). The remaining EAAs (EAA-BCAA) and phenylalanine increased following intake, and more so in the exercising leg at 60 min (p<0.001, Figure 3F and 4D) but there were no differences between groups. Muscle tryptophan decreased similarly in both groups over time, with ∼ 17% lower levels in the exercising leg compared to the resting leg at 180 min (p<0.05, Figure 4F).

### Muscle protein synthesis

Plasma enrichment of L-[*ring*^13^C_6_] phenylalanine was stable during rest, increased slightly during exercise, declined back to resting levels after EAA intake, and thereafter remained relatively stable throughout recovery (Figure 5A). Even though the two groups displayed a highly similar pattern, plasma enrichment was higher in Old compared to Young at multiple timepoints (p<0.05 for all, Figure 5A). Myofibrillar enrichment increased to a greater extent in the exercising leg compared to the resting leg and was generally higher in the Old (p<0.05 for both, Figure 5B). Myofibrillar protein FSR, calculated using plasma enrichment as the precursor, increased in both legs at 60 min (Young, 0.043%·h^-1^ to 0.070%·h^-1^, Old, 0.030%·h^-1^ to 0.070%·h^-1^), with ∼ 50% higher values in the exercising leg compared to the resting leg (p<0.05 for all, Figure 5C). Myofibrillar protein FSR was higher in the exercising leg than the resting leg also at 180 min, however, no differences between Young and Old were observed in any of the timepoints (Figure 5C). Free intracellular enrichment of L-[*ring*^13^C_6_] phenylalanine was elevated compared to basal levels in both legs at 60- and 180 min and these values were higher in Old compared to Young across all timepoints (p<0.05, Figure 5D). When calculated using intracellular enrichment as the precursor, FSR increased by ∼ 95 and 122% after EAA intake and resistance exercise, only in the exercising leg at 60 min and 180 min, respectively (p<0.05 for both, Figure 5E) and was only higher in the exercising leg at 180 min. However, no differences between Young and Old were observed at any time point (p<0.001, Figure 5E).

**Figure 5.**
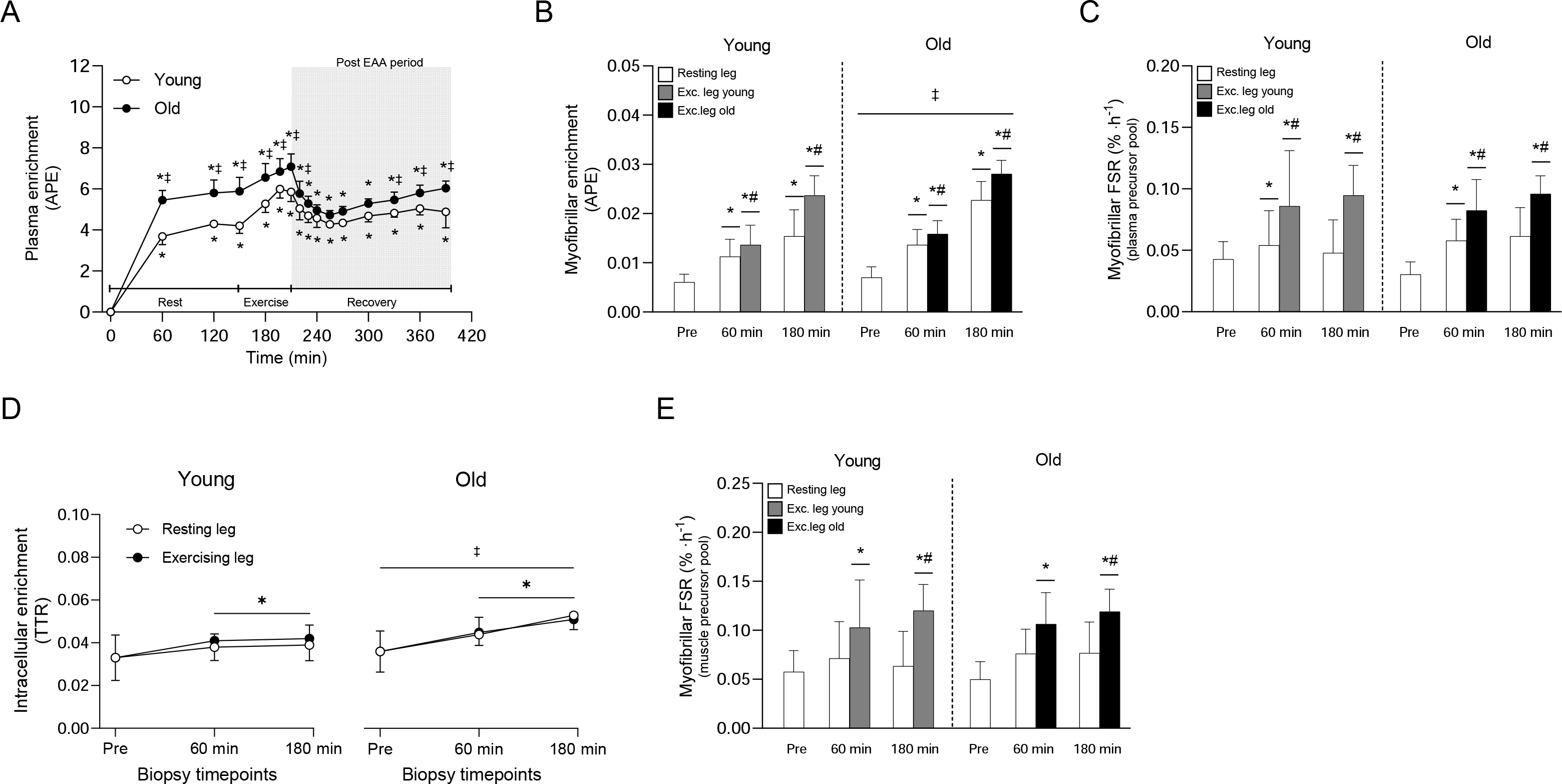
Plasma enrichment of L-[*ring*^13^C_6_] phenylalanine during the experimental trial In Young and Old (A), enrichment of L-[*ring*^13^C_6_] phenylalanine in the myofibrillar fraction in the resting leg and exercising leg of Young and Old (B), myofibrillar protein fractional synthesis rates (FSR) before and after resistance exercise and intake of EAA in the resting and exercising leg of Young and Old determined using plasma enrichment as the precursor (C), free intracellular enrichment of L-[*ring*^13^C_6_] phenylalanine in the resting and exercising leg of Young and Old (D), myofibrillar protein FSR before and after resistance exercise and intake of EAA in the resting and exercising leg of Young and Old determined using free intracellular enrichment as the precursor (E). For plasma variables, white and black circles represent Young and Old, respectively. In muscle, white bars represent the resting leg in both groups, whereas grey and black bars represent the exercising leg in Young and Old, respectively. A two-way (age x time) repeated-measures ANOVA was used to analyzed plasma enrichment and the ANOVA revealed a significant interaction effect (age x time). A three-way (age x time x leg) repeated-measures ANOVA was used to analyze the FSR and the enrichment of the intracellular and myofibrillar fractions. The ANOVA revealed a significant interaction effect (time x leg) for (B, C, E), a significant interaction (age x leg) for (D), a main effect of age for (B) and a main effect of time for (D). The symbols marked without lines represent a three-way interaction, symbols marked with short lines represent a two-way interaction and symbols marked with a long line represent a main effect; * p<0.05 different from baseline, # p<0.05 different from the resting leg, ‡ p<0.05 different from Young. Values are presented as means ± SD for 20 participants.

### Intracellular signaling

In relation to baseline levels, mTOR^Ser2448^ phosphorylation increased following EAA intake and exercise, with a larger increase in the exercising leg compared to the resting leg (∼ 8.2-vs 4.3-fold increase at 60 min, respectively, p<0.001, Figure 6A). In the resting leg, this effect was more pronounced in Old compared to Young at both timepoints (p<0.01 for both, Figure 5A). In a similar fashion, S6K1^Thr389^ and RPS6^Ser240/244^ phosphorylation increased above baseline after EAA intake and exercise, and more so in the exercising leg (p<0.001, Figure 6B and 6C). In the resting leg, phosphorylation was again higher in Old compared to Young at 60- and 180 min (p<0.001 for all, Figure 6B and 6C). Phosphorylation of 4E-BP1^Ser65^ increased following EAA intake and exercise, but the difference between the legs was less pronounced and was statistically different only at 180 min (p<0.001, Figure 6D). However, phosphorylation of this site was on average 78% greater across all timepoints in the Old (p=0.003. eEF2^Thr56^ phosphorylation was reduced (∼ 66 and 39%) in the exercising leg at both timepoints after EAA intake and exercise, with no group differences (p<0.05, Figure 5E). However, phosphorylation at this site was reduced by ∼ 27% also in the resting leg at 60 min, an effect that was only seen in Old (p=0.002, Figure 6E). PRAS40 abundance was on average 38% higher in the Old (p<0.05, data not shown). When related to total levels, PRAS40^Thr246^ phosphorylation increased to the same extent in both groups following EAA intake and exercise and remained elevated throughout the recovery period, with 11-33% higher levels in the exercising leg compared to the resting leg (p=0.003, Figure 6F). Moreover, phosphorylation of all above mentioned proteins, except eEF2^Thr56^, was altered immediately after exercise, with no major differences between groups (p<0.05, supplemental Figure 1).

**Figure 6.**
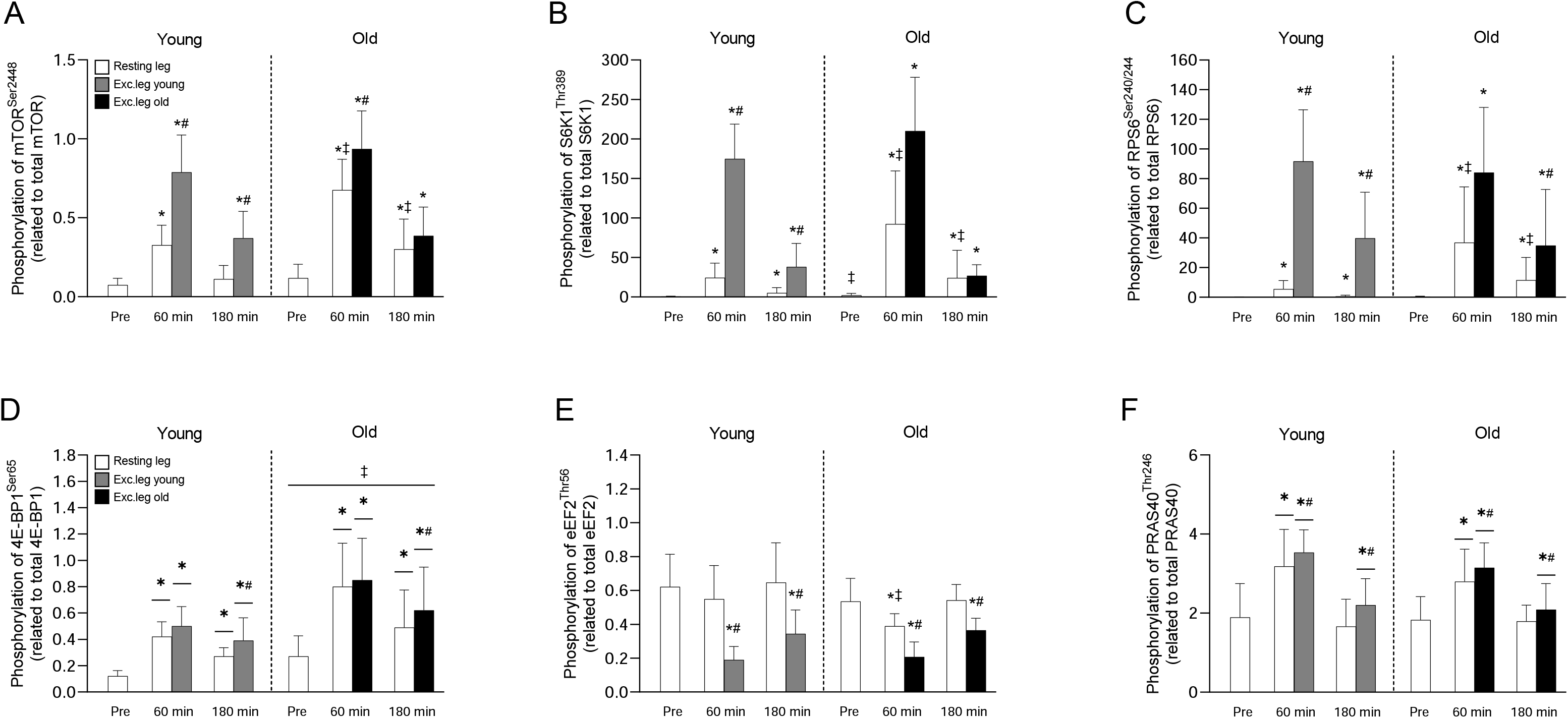
Phosphorylation of mTOR at Ser^2448^ (A), S6K1 at Thr^389^ (B), RPS6 at Ser^240/244^ (C), 4E-BP1 at Ser^65^ (D), eEF2 at Thr^56^ (E) and PRAS40 at Thr^246^ (F) before and after resistance exercise and intake of EAA in Young and Old. White bars represent the resting leg for both groups, whereas grey and black bars represent the exercising leg in Young and Old, respectively. A three-way (age x time x leg) repeated-measures ANOVA was used to analyze phosphorylation state. The ANOVA revealed a significant three-way interaction (age x time x leg) for (A, B, C, E), and a significant two-way interaction (time x leg) for (F and D). There was also a main effect of age for (D). The symbols marked without lines represent a three-way interaction, symbols marked with short lines represent a two-way interaction and symbols marked with a long line represent a main effect; * p<0.05 different from baseline, # p<0.05 different from the resting leg, ‡ p<0.05 different from Young. Values are presented as means ± SD for 20 participants.

In the cytosolic fraction, mTOR content was higher in the exercising leg at 60 min in both Young and Old, but lower in the resting leg at 180 min following EAA intake and exercise only in the Young, compared to baseline (p<0.05 for all, Figure 7A). In the resting leg, mTOR levels were higher in Old compared to Young (p<0.001 for both time points, Figure 7A). In contrast, Rheb and LAT1 levels did not change across time but were approximately 45% and 31% higher, respectively, in the Old at all timepoints (p<0.01 for age, Figure 7B and 7C). In the lysosomal fraction, mTOR content was lower compared to baseline following EAA intake and exercise in the exercising leg at 60 min in both groups (p<0.05, Figure 7D). In addition, Rag B content in the lysosomal fraction was overall ∼ 31% higher in Old compared to Young (p=0.039, Figure 7E). Rag B was also consistently higher in the exercising leg compared to the resting leg (p=0.019, Figure 7E). LAMP2 levels did not change over time or between groups (Figure 7F).

**Figure 7.**
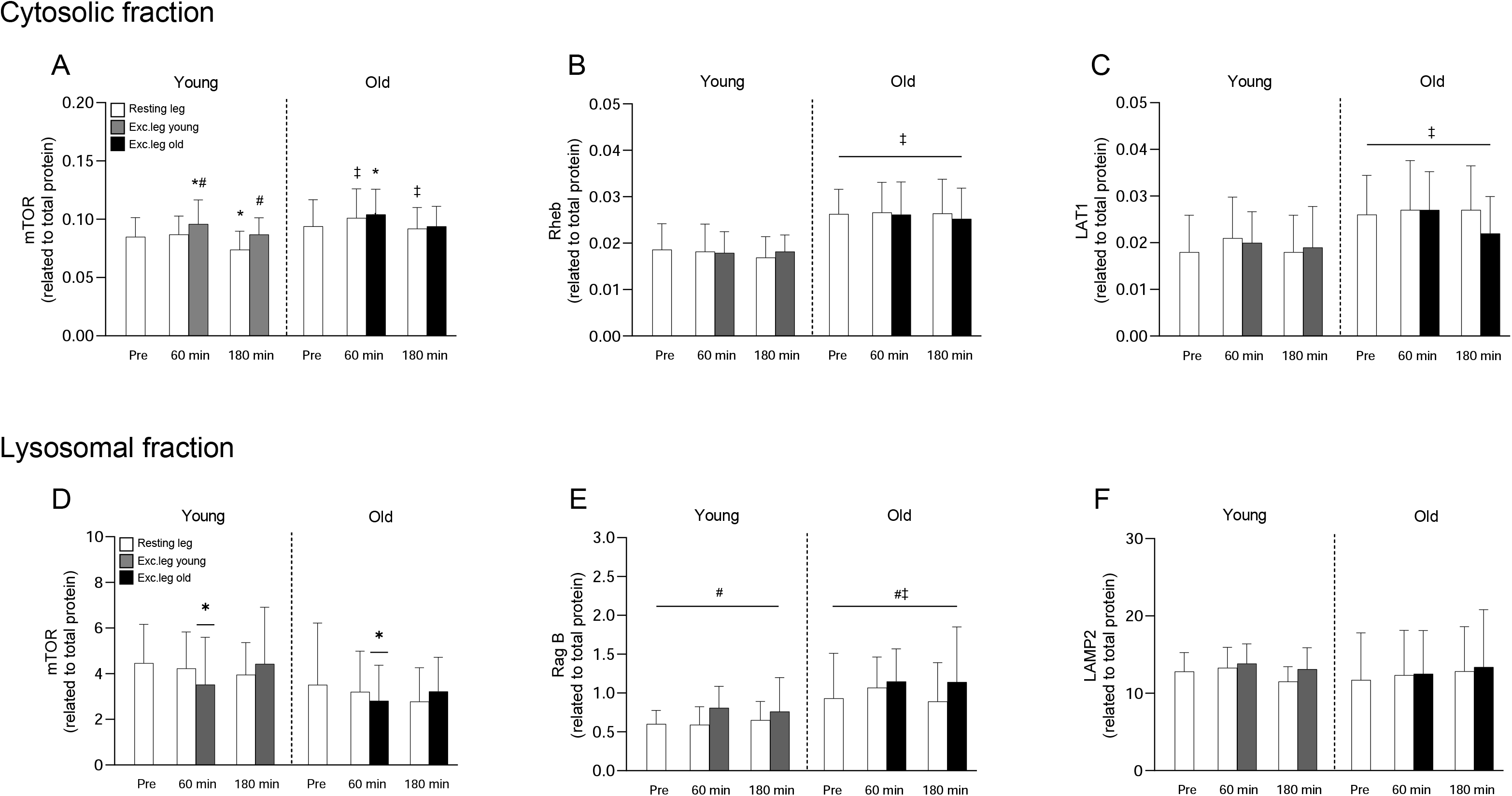
Total levels mTOR (A), Rheb (B), and LAT1 (C) in the cytosolic fraction and total levels of mTOR (D), Rag B (E) and LAMP2 (F) in the lysosomal fraction before and after resistance exercise and intake of EAA in Young and Old. White bars represent the resting leg in both groups, whereas grey and black bars represent the exercising leg in Young and Old, respectively. A three-way (age x time x leg) repeated-measures ANOVA was used to analyze the data. The ANOVA revealed a significant interaction effect (age x time x leg) for (A) and a main effect of age for (B, C). Furthermore, the ANOVA revealed a significant interaction effect (time x leg) for (D) and main effects of age as well as leg for (E). The symbols marked without lines represent a three-way interaction, symbols marked with short lines represent a two-way interaction and symbols marked with a long line represents a main effect; * p<0.05 different from baseline, # p<0.05 different from the resting leg, ‡ p<0.05 different from Young. Values are presented as means ± SD for 20 participants.

## Discussion

Currently, the prevailing view is that skeletal muscle of old individuals has a reduced ability to stimulate muscle protein synthesis (MPS) in response to anabolic stimuli, and this anabolic resistance (AR), is believed to be the primary cause of sarcopenia [3, 4]. However, the presence of AR in old skeletal muscle is not a universal finding [13]. The reason for these inconsistencies is not readily apparent but may be related to variables secondary to old age. For instance, ageing is often accompanied by increased adiposity and reduced physical activity [25, 26], both of which have been shown to independently induce AR [14–17]. Thus, AR may be mediated by diet and levels of physical activity in older adults, rather than by ageing alone. To test this question, we assessed AR in lean, healthy and physically active young and old men. In contrast to our initial hypothesis, when controlling for adiposity and habitual physical activity, we found that AR was not present, with the anabolic response to amino acid intake, both alone and in combination with resistance exercise, maintained in old skeletal muscle.

Several physiological mechanisms have been proposed as contributing factors to the development of AR, including increased splanchnic extraction of exogenous amino acids, particularly leucine [27, 28]. Here, we found that plasma amino acids peaked at similar levels in both groups (Figure 3 and 4), indicating that absorption and subsequent release of amino acids into circulation is not diminished with increasing age. However, despite similar peak levels, plasma clearance was more rapid during the recovery phase in young, an effect which was more pronounced for the branched-chain amino acids (BCAA) than the remaining EAA. On its own, this could indicate an increased incorporation into new proteins through increased MPS but given the lack of difference in MPS between the two groups, it is more likely a result of a larger distribution volume in the young.

After intake, and in absolute terms, muscle levels of EAA increased to the same extent in both groups, whereas tryptophan, which was not provided orally, gradually decreased, see Figure 4. This effect was more pronounced in the exercising leg, likely because of higher rates of MPS. Interestingly, when expressed as fold-change, the old had a larger increase in muscle amino acid levels, particularly of the BCAAs (Supplemental figure 1). These findings are in line with those of Cuthbertson et al. who found higher leucine levels in anabolically resistant old men. They concluded that the increased leucine levels were likely a consequence of lower incorporation into muscle proteins as evidenced by the blunted MPS response [11]. However, the similar MPS response between young and old seen here, suggests that the elevated amino acids levels in the old may be a result of increased uptake rather than decreased incorporation. This may indeed be the case as we found approximately 30% greater abundance of the amino acid transporter LAT1 (SLC75A) in the old group (Figure 7), which is primarily responsible for transporting large neutral amino acids, including leucine, into the cell [29, 30]. These data are partly in line with work showing that older adults increase LAT1 protein levels in response to mechanical loading, possibly to prime the cell for increased amino acid availability [31]. However, rather than being changed dynamically with muscle contractions, our data indicate that LAT1 is upregulated in aged muscle independent of acute exercise. While the reason for this remains to be fully determined, it may reflect an adaptive cellular mechanism to ensure proper transport during periods of low amino acid availability or to offset a potential deficit in other amino acid transporters, i.e., LAT2 (SLC7A8).

At the cellular level, MPS is largely regulated by the mTORC1 signalling pathway [5, 6]. As such, it is reasonable to assume that the AR observed in several studies, would be a consequence of dysregulated mTORC1 signalling. While there is indeed some support for this notion [11, 12], the upstream mechanisms responsible for such dysfunction remain unclear. Mechanistically, lysosomal/mTORC1 movement towards the cell periphery [32], and increased translocation of mTORC1 to the lysosomal surface in response to amino acid provision *in vitro* [9, 10], have both been implicated in the activation of the complex. As such, alterations in these cellular events may represent key mechanisms underlying AR. Shuttling of mTORC1 to the lysosomal surface is largely regulated by the RAG GTPases. In the presence of amino acids, RagB forms a complex with either RagC or RagD, and subsequently recruits mTORC1 to the lysosomal surface where it becomes fully activated by Rheb [9, 10]. Interestingly, here we found that both RagB and Rheb were markedly enriched in aged muscle despite no differences in lysosomal content, as assessed by the lysosomal marker LAMP2, see Figure 7. The biological relevance of having greater basal pools of these proteins is not readily apparent as this, to the best of our knowledge, has not been described previously in the context of skeletal muscle. However, RagB serves as an integral part of the amino acid sensing machinery [9], and in other cell types such as neurons which are fundamental for organismal survival, increased levels of Rag B constitute a mechanism for keeping mTORC1 active in conditions of low amino acid availability [33]. Furthermore, as the most proximal activator of mTORC1, Rheb is essential for the kinase activity of the complex [34], and overexpression of Rheb has been shown to induce muscle hypertrophy [35]. Together with the increased levels of LAT1, our findings point toward a compensatory but coordinated response aimed at maintaining functionality of several key cellular events required to maintain the anabolic response of aged muscle.

In addition to increased targeting to the lysosomal surface [9, 10], trafficking of mTORC1 towards the cell periphery appears to be crucial for activation of the complex [32], a finding also observed in multiple studies in human skeletal muscle following resistance exercise and protein feeding [18, 36–38]. In contrast, human muscle data on feeding and exercise-induced changes in lysosomal/mTORC1 interactions are less clear, with studies showing increased [19, 36, 39], decreased [20, 37, 38] as well as unaltered [18], interactions, as assessed by mTOR and LAMP2 co-localization. Here, we were unable to measure lysosomal/mTORC1 trafficking, but could demonstrate a lower abundance of mTOR in the lysosomal fraction, indicative of a decreased interaction with the lysosome, at the 60-min timepoint in the exercising leg (Figure 7). Our findings are thus in line with previous studies showing a decreased lysosomal/mTORC1 interaction following exercise and feeding [20, 37, 38]. Interestingly, this coincided with a parallel increase of mTOR abundance in the cytosolic fraction and the most robust phosphorylation of S6K1 and its downstream target rpS6. This suggests that the cytosol is the major site for mTORC1-dependent phosphorylation of S6K1, as previously shown in non-muscle tissue [40]. Here, this notion is supported by S6K1 primarily being located in the cytosolic fraction (data not shown). Whether or not these phosphorylation events occurred closer to the cell periphery in the present study is unclear. However, it appears likely as the cell periphery is the cellular region at which phosphorylation of rpS6 is most potently increased following exercise and feeding [41]. Regardless, there were no obvious differences in mTORC1 abundance between young and old in the two fractions, which is in line with the similar phosphorylation patterns observed in the two groups, see Figure 6.

Interestingly, the robust differences in protein content of ∼30-45% between groups did not translate into overall differences in MPS or mTORC1 signalling (Figure 5 and 6). This indicates that the upregulation of amino acid signalling seen here may be a compensatory mechanism in old skeletal muscle. This compensation does not appear to be related to insulin signalling or peptide elongation as we observed similar changes in PRAS40 ^Thr246^ and eEF2^Thr56^ phosphorylation, respectively (Figure 6). Furthermore, while there appears to be a causal relationship between upregulated protein content and the maintained anabolic response in the old cohort, it remains unclear whether these changes represent an adaptive response aimed at delaying age-induced muscle loss, or if it reflects the beginning of a dysfunctional mTORC1 pathway, as shown in some studies where mTORC1 activity is upregulated in the basal state of aged muscle [42, 43]. Here, we did find slightly but significantly higher S6K1^Thr389^ phosphorylation in the old group at baseline, see Figure 6. However, the physiological relevance of this minute difference is unclear as the capacity to increase MPS after resistance exercise and EAA intake was highly preserved in the old group. Regardless, our findings clearly show that the anabolic potential is well preserved in healthy, lean aged human skeletal muscle.

From a physiological perspective, several potential explanations may underline the findings presented here. As mentioned previously, ageing is often associated with increased adiposity [25], which in itself can induce AR [15], and has also been shown to be predictive of muscle loss with old age [44]. In several of the previous studies, the participants were indeed overweight (BMI 26-27) [11, 12, 45], which may have been a contributing factor to the observed AR. In addition to adiposity, physical activity levels also play an important role in maintaining the anabolic potential of muscle [14, 16, 17]. Here, the participants in the old cohort were recreationally active 2-3 times per week. As such, the relatively high level of physical activity, by itself or in concert with the low BMI (<23), may have contributed to maintaining anabolic sensitivity in the present study. Additional contributing factors may include the design of the exercise intervention. Here we chose an exercise regimen based each participant’s 10 RM in which each set was to be performed very close to volitional failure. Despite the lower absolute load in the old cohort, the high volume together with the high degree of metabolic stress likely provided a maximal stimulus, thus facilitating the same anabolic response in both groups. This would be in line with previous research showing that in young individuals, low load, but high-volume resistance exercise performed to exhaustion, is highly potent in stimulating MPS [46].

Taken together, here we demonstrated that lean, healthy and physically active older adults do not manifest anabolic resistance when performing high-intensity high volume resistance exercise in combination with EAA intake. The maintained anabolic sensitivity of the old group appears to be underpinned by increased expression of proteins involved in amino acid transport (LAT1), amino acid sensing (RagB) and activation (Rheb) of mTORC1. These findings may represent a compensatory mechanism of healthy aging muscle to retain anabolic sensitivity, potentially to delay the inevitable muscle loss ultimately seen with very old age. To minimize confounding factors, future studies examining the role of old age *per se* in AR should consider the adiposity and physical activity levels of the participants and use exercise interventions that elicit maximal protein synthetic responses.

## Figure legends

**Supplemental figure 1.**
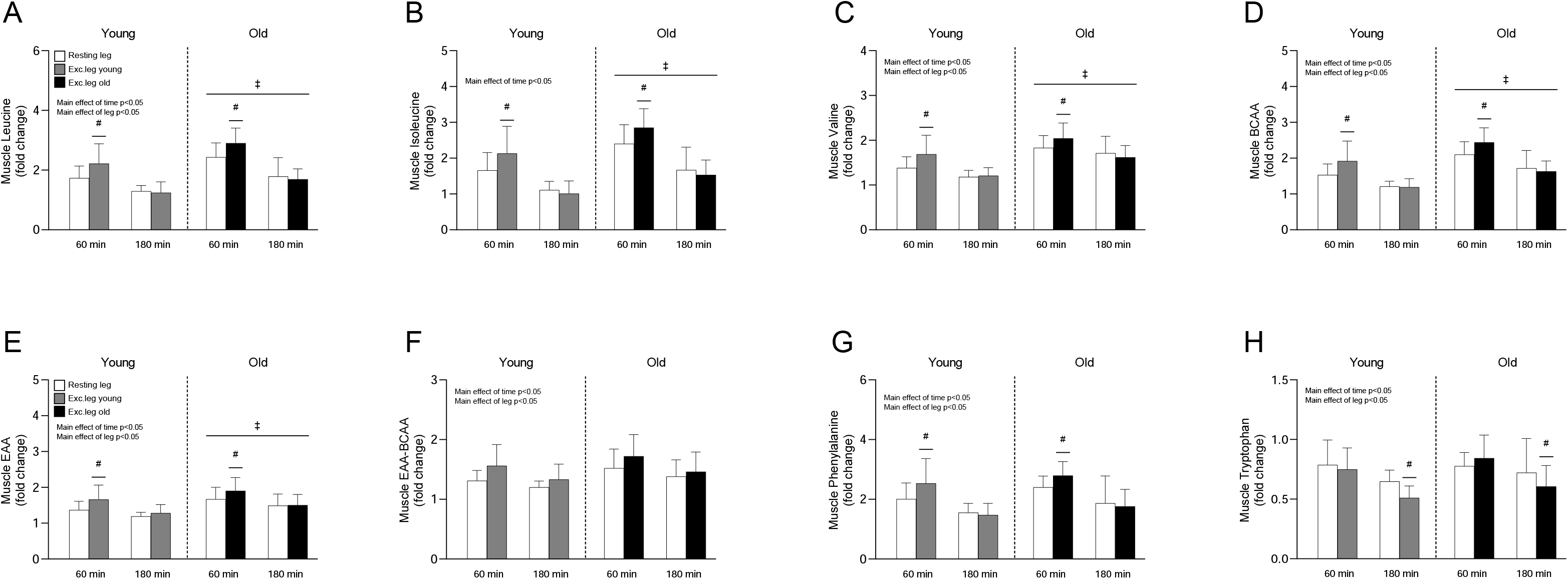
Muscle concentrations of leucine (A), isoleucine (B), valine (C), BCAA (D), EAA (E), EAA-BCAA (F), phenylalanine (G) and tryptophan (H) presented as fold changes before and after resistance exercise and intake of EAA in Young and Old. White bars represent the resting leg in both groups, whereas grey and black bars represent the exercising leg in Young and Old, respectively. A three-way (age x time x leg) repeated-measures ANOVA was used to analyze the data. The ANOVA revealed a significant interaction effect (time x leg) for (A, B, C, D, E, G, H), main effects of age for (A, B, C, D, E) and main effects of leg for (A, B, C, D, E, G, H). The symbols marked with short lines represent a two-way interaction and symbols marked with a long line represent a main effect; * p<0.05 different from baseline, # p<0.05 different from the resting leg, ‡ p<0.05 different from Young. Values are presented as means ± SD for 20 participants.

**Supplemental figure 2.**
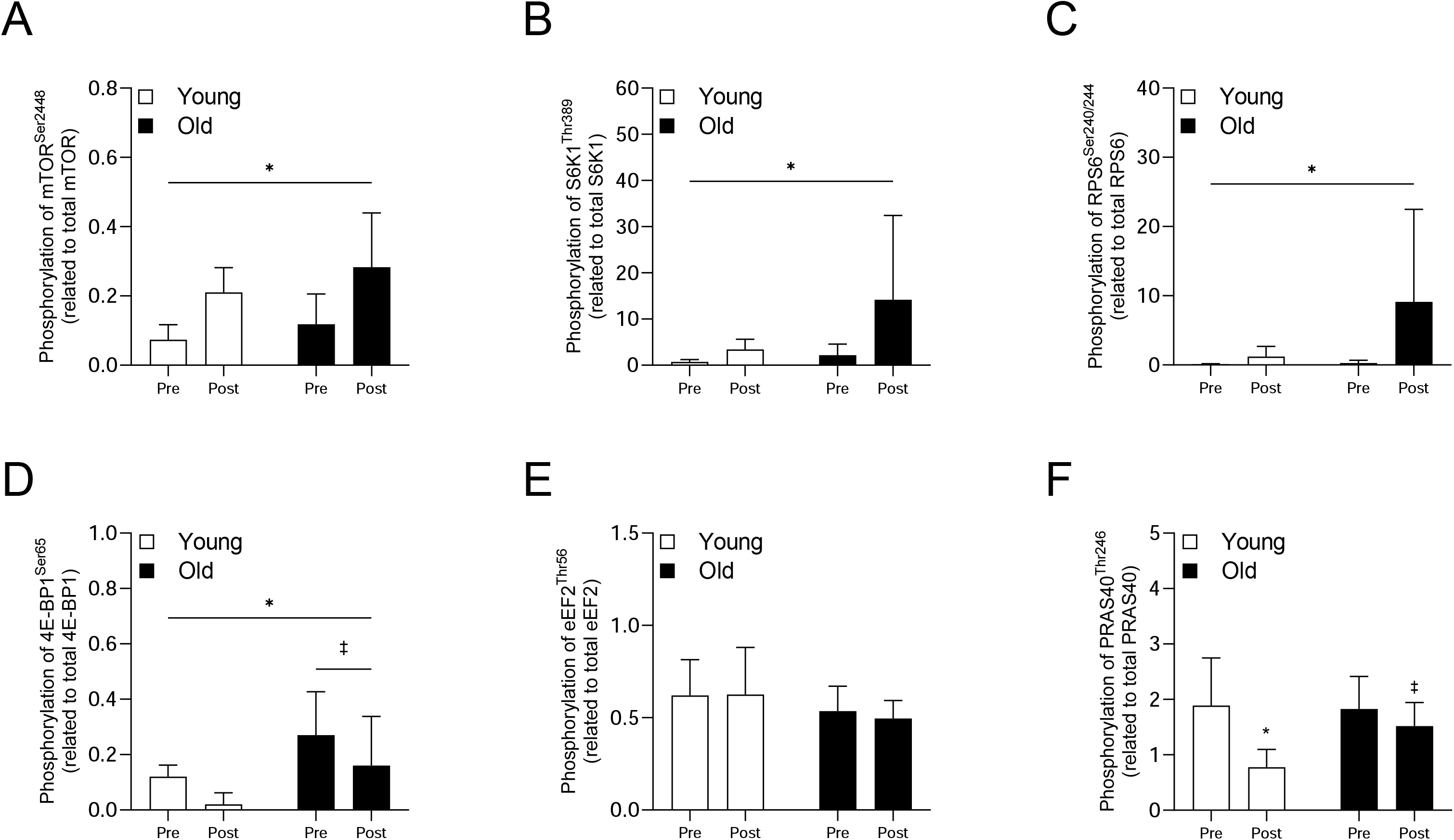
Phosphorylation of mTOR at Ser^2448^ (A), S6K1 at Thr^389^ (B), RPS6 at Ser^240/244^ (C), 4E-BP1 at Ser^65^ (D), eEF2 at Thr^56^ (E) and PRAS40 at Thr^246^ (F) before and immediately after resistance exercise in Young (white bars) and Old (black bars). A two-way (age x time) repeated-measures ANOVA was used to analyze the data. The ANOVA revealed a significant interaction effect (age x time) for (F), a main effect of time for (A, B, C, D) and a main effect of age for (D). The symbols marked without lines represent a two-way interaction and symbols marked with lines represent a main effect. * p<0.05 different from baseline or main effect of time, ‡ p<0.05 different from Young. Values are presented as means ± SD for 20 participants.

**Supplemental figure 3.**
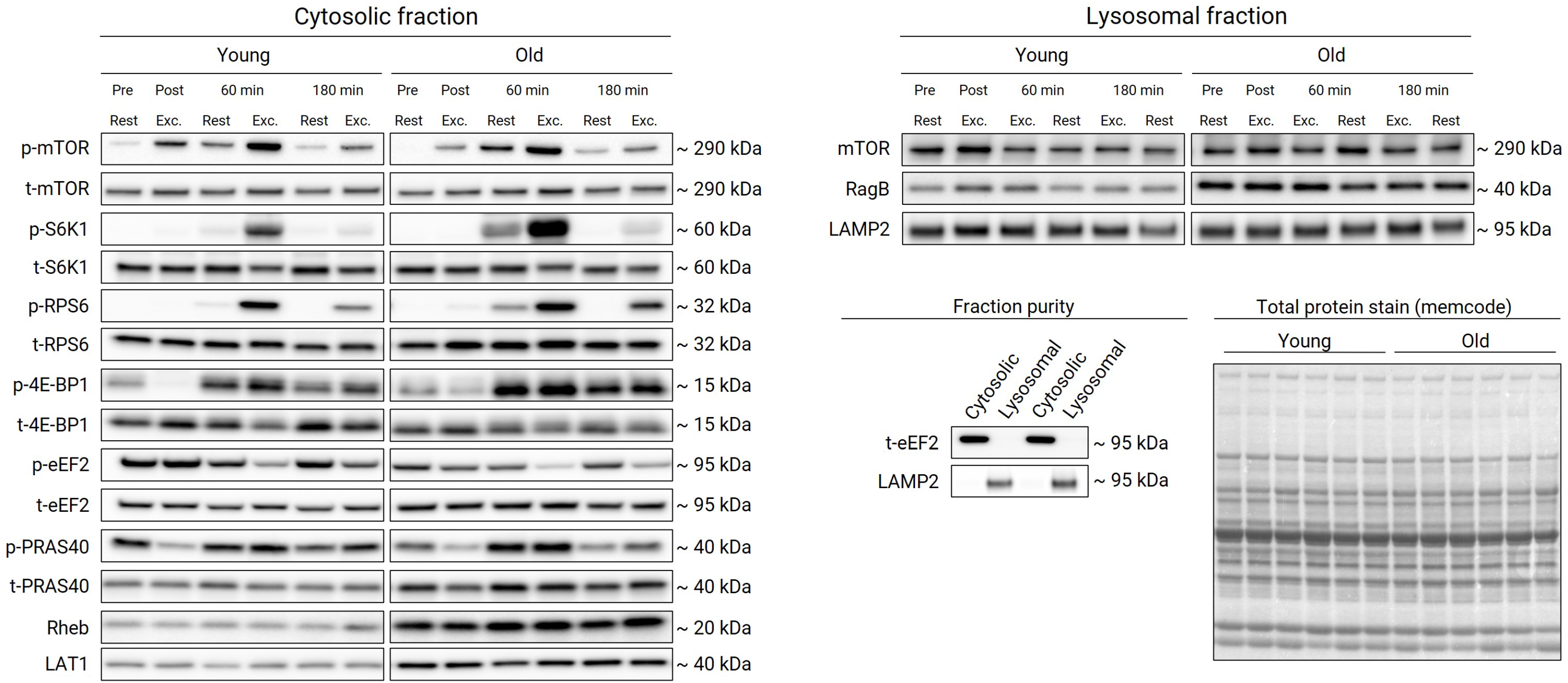
Representative images of the immunoblots for the proteins presented in Figure 6 and 7. The left panel illustrates the phosphorylated and total forms of proteins residing in the cytosolic fraction and the right panel illustrates proteins residing in the lysosomal fraction. Representative blots are presented from one young and one old participant. In the bottom right panels, the purity of the two different fractions as well as the total protein stains (MemCode™ Reversible Protein Stain) are shown.

## Funding

This project was supported by funding from the European Research Council (ERC) under the European Union’s Horizon 2020 research and innovation programme (grant agreement #707336) to AP, The Åke Wiberg Foundation (grant #M17-0259) and The Lars Hiertas Memorial Foundation (grant# FO2017-0325) to WA, and the Elisabeth and Gunnar Liljedahl Foundation to OH.

## Acknowledgements

The authors wish to thank the participants for their time and effort. The authors also wish to thank Prof. Leigh Breen and Prof. Gareth Wallis for assistance during sample collection, Prof. Björn Ekblom and Dr Yasir S Elhassan for medical supervision, and Prof. Philip J Atherton and Prof. Stephen D R Harridge for fellowship supervision.

## Conflict of interest

The authors declare that they have no conflicts of interest.

## Notes

### Competing Interest Statement

The authors have declared no competing interest.

